# Synthetically Engineered *Medea* Gene Drive System in the Worldwide Crop Pest, *D. suzukii*

**DOI:** 10.1101/162255

**Authors:** Anna Buchman, John M. Marshall, Dennis Ostrovski, Ting Yang, Omar S. Akbari

## Abstract

Synthetic gene drive systems possess enormous potential to replace, alter, or suppress wild populations of significant disease vectors and crop pests; however, their utility in diverse populations remains to be demonstrated. Here, we report the creation of the first-ever synthetic *Medea* gene drive element in a major worldwide crop pest, *D. suzukii*. We demonstrate that this drive element, based on an engineered maternal “toxin” coupled with a linked embryonic “antidote,” is capable of biasing Mendelian inheritance rates with up to 100% efficiency. However, we find that drive resistance, resulting from naturally occurring genetic variation and associated fitness costs, can hinder the spread of such an element. Despite this, our results suggest that this element could maintain itself at high frequencies in a wild population, and spread to fixation, if either its fitness costs or toxin resistance were reduced, providing a clear path forward for developing future such systems.

## Introduction

Spotted wing *Drosophila, D. suzukii*, is a major worldwide crop pest of various soft-skinned fruits ^1^. Unlike other *Drosophilids* that prefer to oviposit on overripe fruits, *D. suzukii* utilizes its serrated ovipositor to lay eggs inside ripening fruits, causing significant crop losses ^1-3^. Found only in Japan prior to the 1930’s ^4^, in the last several decades *D. suzukii* has spread invasively to every continent except Antarctica ^1,5^. In the United States, for example, *D. suzukii* was initially discovered in Santa Cruz, California, in 2008, and since then has rapidly invaded many states and is a significant threat to fruit industries across the country ^2^. For example, between 2009-2014, *D. suzukii* caused an estimated $39.8 million in revenue losses for the California raspberry industry alone ^6^, and is responsible for 20%-80% crop losses in other fruit production areas ^1,3,4,6^. Current methods to control *D. suzukii* rely considerably on the use of expensive, broad-spectrum insecticides (e.g., malathion), which have variable efficacy ^5^, are difficult to use due to timing of fruit infestation ^7^, and face the risk of *D. suzukii* evolving resistance ^8^. While other forms of control may be possible (e.g., the use of recently identified natural predators ^9^ or oral delivery of dsRNA by microbes ^10^), these approaches have not been widely adopted ^11,12^. Therefore, given the rapid worldwide spread and potential economic impact of *D. suzukii*, novel effective control measures are urgently needed.

An alternative approach that would complement existing control methods would be the use of engineered *D. suzukii* as a genetic-based control strategy ^13^. Use of genetically modified insects for wild population manipulation was first suggested over a half century ago ^14–16^, and has garnered considerable interest in recent years ^17–19^. In fact, one method of using genetically modified insects for population control, a system called RIDL (Release of Insects carrying a Dominant Lethal) ^20–22^, where males carrying a repressible dominant lethal transgene are released to mate with wild females and produce non-viable progeny, has recently been implemented in the field. Although this strategy has been shown to be effective in reducing insect populations ^23–25^, it requires continuous rearing and ongoing inundative releases of large numbers of individuals, making it rather costly and labor-intensive; furthermore, it has not been developed for *D. suzukii*.

Other proposed methods of using genetically modified organisms for population control rely on engineered gene drive systems that function by forcing their inheritance in a non-Mendelian fashion, allowing the drives to catalytically increase in frequency with each generation even without conferring fitness advantages to their host ^17,21,26^. Such methods could be utilized to spread desirable genes through populations or even to suppress target populations 27, and are promising self-sustaining tools for various applications where manipulation of wild populations may be desirable ^18,28^. A number of engineered gene drive mechanisms have been proposed ^17,18,21,26,27,29^; however, to date, only *Medea* (Maternal Effect Dominant Embryonic Arrest) and an underdominance-based approach have been demonstrated to bring about robust population replacement in wildtype laboratory populations ^30–32^. Specifically, *Medea* systems rely on expression of a toxin-antidote combination, such as a microRNA (miRNA) toxin that is expressed during oogenesis in *Medea*-bearing mothers, and a tightly linked antidote expressed early during embryogenesis in *Medea*-bearing progeny (Figure 1A). The toxin is inherited by all progeny from a *Medea*-bearing mother, resulting in miRNA-mediated suppression of an essential embryonic gene that causes disruption of normal development during embryogenesis (Figure 1A-C). Offspring that inherit *Medea* receive a tightly linked antidote, consisting of a zygotically active miRNA-resistant copy of the targeted essential gene, that allows for restoration of normal development (Figure 1D); non-*Medea*-bearing progeny from *Medea*-bearing mothers lack this antidote and perish (Figure 1E). Due to this dominant inheritance, *Medea* is predicted to rapidly spread itself, and any linked cargo genes, through a target population ^30–32^.

**Figure 1.**
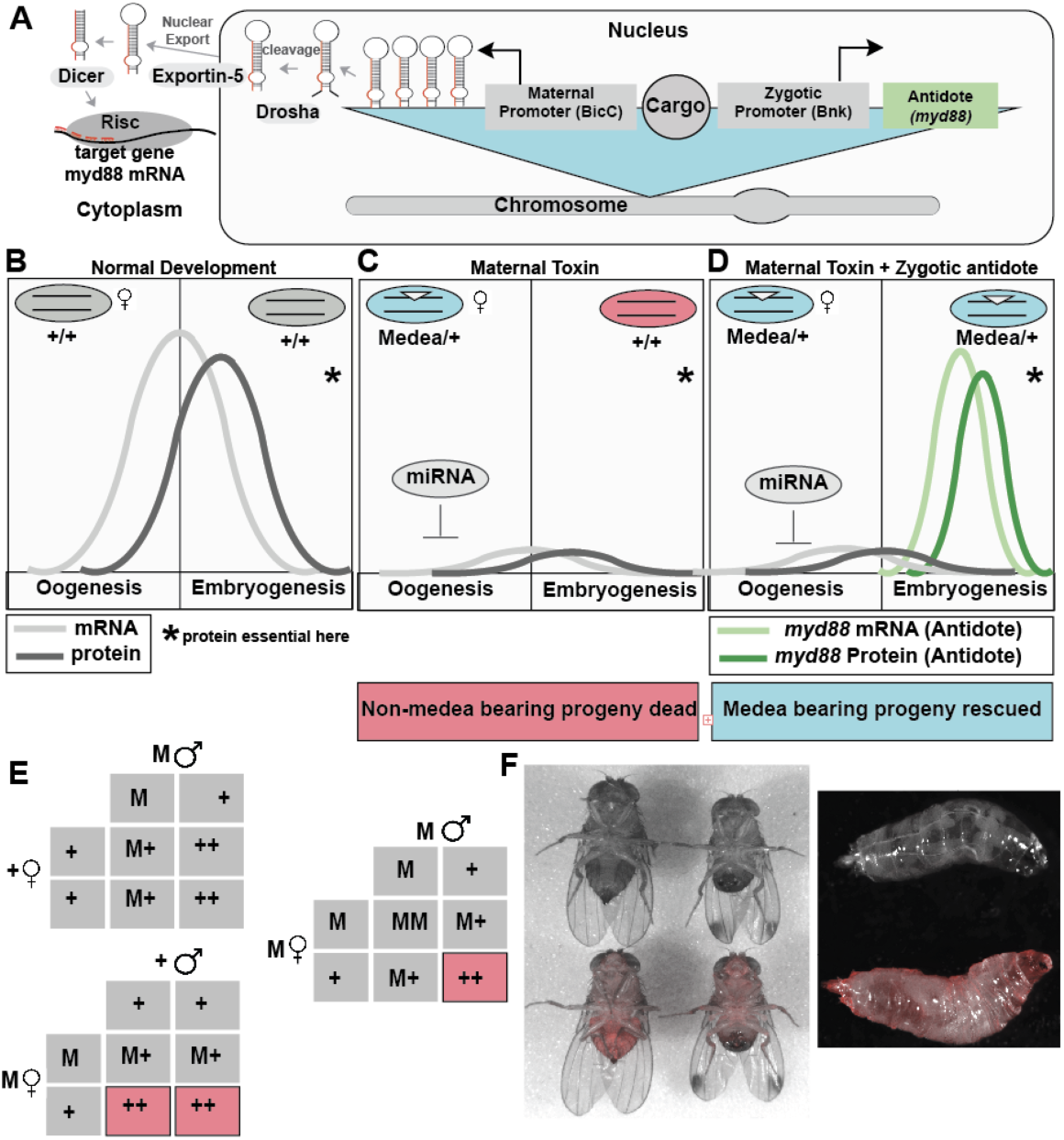
A synthetic *Medea* element in *D. suzukii*. *D. suzukii Medea* transgene was generated to comprise a miRNA “toxin” targeting the 5’ UTR of *D. suzukii* myd88 expressed under the predicted *D. suzukii* female germ-line–specific bicoid (BicC) promoter, an “antidote” consisting of *D. suzukii* myd88 coding region driven by the predicted *D. suzukii* early embryo–specific bottleneck (bnk) promoter, and two separate transformation markers – eGFP under control of the eye-specific 3xP3 promoter, and dsRed under control of the ubiquitous hr5-IE1 promoter (A). During normal development maternal myd88 deposited into the embryo, where it is required for normal development (B). The *Medea* miRNA toxin targets myd88 mRNA during oogenesis, preventing proper deposition into the embryo and causing embryonic lethality in progeny that lack the *Medea* element (C). In embryos that possess a copy of the Medea element, a version of myd88 that is insensitive to the miRNA toxin is expressed during early embryogenesis, rescuing miRNA-induced lethality (D). When heterozygous Medea males are crossed out to wild type females, all progeny survive since the maternal toxin is not expressed; however, when heterozygous *Medea* females are crossed to wild type males, 50% of the progeny - the ones that fail to inherit *Medea* perish. When heterozygous females are crossed to heterozygous males, 75% of the progeny inherit *Medea*, either from the mother or the father, and survive, while those that fail to inherit a *Medea* element perish (E). The hr5-IE1 promotes robust expression of dsRed in both D. suzukii adults and larvae, allowing for facile identification of Medea-bearing individuals (F).

For *D. suzukii*, since suppression of the pest population is ultimately desired, a synthetic *Medea* could be used to achieve suppression by spreading a cargo gene proffering susceptibility to a particular chemical, or by driving in a conditional lethal gene activated by an environmental cue such as diapause ^32^. Therefore, given the potential utility of a *Medea* system in *D. suzukii*, we leveraged the limited genetic tools and techniques available in this non-model organism, for example the draft genome assembly ^33^ and transgenesis ^34^, to engineer the first *Medea*-based population control technology. As *D. suzukii* is generally poorly genetically characterized, we had to overcome numerous challenges to accomplish this feat, e.g., identifying and testing necessary components such as maternal and zygotic promoters and demonstrating the ability to engineer miRNAs that target desired sequences. Notwithstanding, we overcame these challenges, and herein describe the successful development of a potent *Medea* system in *D. suzukii*. We demonstrate that this system is capable of drastic dominant inheritance to achieve non-Mendelian transmission frequencies of up to 100% in many geographically distinct populations. This represents, to our knowledge, the first proof-of-principle gene drive mechanism developed in a crop pest.

## Results

### Generation of Synthetic *Medea* Element

To create a synthetic *Medea* element in *D. suzukii*, we engineered a *PiggyBac* vector comprising a miRNA toxin coupled with a toxin-resistant antidote, inspired by the architectures used to generate previous *Medea* systems in *D. melanogaster* ^32,35^. We designed synthetic miRNAs to target *D. suzukii* myd88, a highly conserved gene shown to be maternally deposited and required for dorsal-ventral patterning in the early embryo in *D. melanogaster* ^36^. We used the predicted *D. suzukii* female germline-specific bicoid (BicC) promoter to drive expression of a “toxin” consisting of a polycistronic set of four synthetic microRNAs (miRNAs) each designed to target the 5’ untranslated region (UTR) of *D. suzukii* myd88 (Figure 1A). Importantly, to ensure these miRNAs could target the desired sequence, we performed genomic DNA sequencing of the myd88 5’UTR target region in our reference *D. suzukii* strain (collected from Corvallis, Oregon) and designed the miRNAs against this sequence (Supplementary Figure 1).This *Medea* element also contained an “antidote” consisting of the *D. suzukii* myd88 coding region, insensitive to the miRNAs as it does not contain the miRNA-targeted 5’UTR, driven by the predicted *D. suzukii* early embryo-specific bottleneck (bnk) promoter, and two separate transformation markers – eGFP driven by the eye-specific 3xP3 promoter ^37^, and dsRed driven by the ubiquitous hr5-IE1 promoter ^38^.

### Characterization of *Medea* Genetic Behavior

Following microinjection of the *Medea* transgene into *D. suzukii* embryos, a single G_1_ transformant male was recovered, as identified by ubiquitous hr5-IE1 driven expression of dsRed (Figure 1F), and weak eye-specific 3xP3-driven eGFP. When outcrossed to several wildtype (non-*Medea* bearing; +/+) females, this male produced roughly ∼50% *Medea-*bearing and ∼50% wildtype offspring, as would be expected from standard Mendelian segregation without dominance (Table 1). Resulting heterozygous G_2_ *Medea*-bearing progeny were individually outcrossed to wildtype individuals of the opposite sex to determine inheritance patterns, and these individual outcrosses were continued for six generations (Table 1). Remarkably, until the G_5_ generation, all heterozygous *Medea*/+ mothers (n = 91) produced 100% *Medea*-bearing progeny (n = 1028), while heterozygous *Medea*/+ fathers (n = 16) produced ∼50% *Medea*-bearing progeny (n = 268). While the majority of heterozygous *Medea*/+ G_5_ (23/31) and G_6_ (16/25) generation females also produced 100% *Medea*-bearing progeny, some heterozygous G_5_ (8/31), and G_6_ (9/25) females unexpectedly produced a small yet notable number (52/1219) of wildtype offspring. Notwithstanding, individually these G_5_ and G_6_ heterozygous *Medea*/+ females displayed significantly biased dominant inheritance rates ranging from 76%-96%, with an average rate of 86.4%. Overall, in six generations of individual female outcrosses, the percentage of *Medea*-bearing progeny borne by single heterozygous *Medea*/+ mothers (n = 147) was 97.7% (2195/2247; Table 1) as opposed to the 50% that would be expected with standard Mendelian segregation without dominance, indicating that the *Medea* element is extremely functional at biasing inheritance.

**Table 1.** *D. suzukii Medea* shows predicted genetic behavior. Results of heterozygous *Medea D. suzukii* individual fly outcrosses to wild type *D. suzukii*. G_1_ indicates the offspring from injected G_0_ individuals, with subsequent numbers (G_2_-G_6_) indicating subsequent generations.

| Generation | Sex (# crossed) | # of Progeny | Average % *Medea* |
| --- | --- | --- | --- |
| G<sub>1</sub> | ♂ (1) | 22 | 54.5% |
| G<sub>2</sub> | ♀ (9) | 126 | 100% |
| G<sub>2</sub> | ♂ (3) | 45 | 43.3% |
| G<sub>3</sub> | ♀ (32) | 299 | 100% |
| G<sub>3</sub> | ♂ (12) | 201 | 48.8% |
| G<sub>4</sub> | ♀ (50) | 603 | 100% |
| G<sub>5</sub> | ♀ (31) | 785 | 96.8% |
| G<sub>6</sub> | ♀ (25) | 434 | 93.8% |
| ***Medea*+/total individuals from females (147)** |  | **2195/2247** | **97.7%** |

### *D. Suzukii Medea* Exhibits Maternal-Effect Lethality and Zygotic Rescue

To further characterize the genetics behind the highly biased inheritance patterns described above, additional crosses between individuals of various *Medea* genotypes were performed, and confirmed that *Medea* exhibits maternal-effect lethality and zygotic rescue (Table 2). For example, matings between heterozygous *Medea*/+ mothers and wildtype fathers resulted in 55.63±0.76% embryo survival with 94.20±1.33% of the progeny being *Medea*- bearing, while matings between heterozygous *Medea/*+ mothers and heterozygous *Medea/*+ fathers yielded 79.11±3.95% embryo survival with 94.12±0.67% of the progeny being *Medea*- bearing. The higher-than-expected embryo survival is consistent with the observation that not all heterozygous *Medea*/+ mothers give rise to 100% *Medea*-bearing progeny, indicating that not all wildtype progeny from a heterozygous *Medea*/+ mother perish.

**Table 2.** *D. suzukii Medea* chromosomes show maternal-effect lethality and zygotic rescue. Crosses between parents of specific genotypes (indicated in the two leftmost columns) were carried out, and progeny survival to crawling first-instar larvae was quantified (third column from right). M indicates *Medea*, + indicates wild-type, red text indicates genotypes expected to be inviable. The percentage of transgenic adults resulting from each cross type was quantified (rightmost column).

| Parental Genotype |  | Progeny Genotype (%) | Embryo Survival % |  | Adult Transgenic % |  |
| --- | --- | --- | --- | --- | --- | --- |
| ♀ | ♂ |  | Predicted | Observed | Predicted | Observed |
| M/M | M/M | M/M (100%) | 100 | 97.63±0.25 | 100 | 100 |
| M/M | +/+ | M/+ (100%) | 100 | 95.67±1.87 | 100 | 100 |
| +/+ | M/M | M/+ (100%) | 100 | 97.07±0.42 | 100 | 100 |
| M/+ | M/+ | M/M (25%)<br>M/+ (50%)<br>+/+ (25%) | 75 | 79.11±3.95 | 100 | 94.12±0.67 |
| M/+ | +/+ | M/+ (50%)<br>+/+ (50%) | 50 | 55.63±0.76 | 100 | 94.20±1.33 |
| +/+ | +/M | M/+ (50%)<br>+/+ (50%) | 100 | 89.74±1.39 | 50 | 48.39±3.22 |

### *Medea* Functionality in Geographically Distinct Populations

To assess whether the *D.suzukii Medea* could function in geographically distinct populations especially likely to harbor genetic variability in regions that canonically have less conservation such as the 5’UTR, heterozygous *Medea*/+ flies were tested in eight additional *D. suzukii* strain backgrounds. These strains were collected from various locations around the world, including: Mt. Hood, OR; Clayton, WA; Brentwood, CA; Tracy, CA; Watsonville, CA; Oahu, HI; Beltsville, MD; and Ehime, Japan. Interestingly, for 3/8 strains, the *Medea* inheritance rate from heterozygous *Medea*/+ mothers was 100%, while from 5/9 strains the inheritance rate ranged from 87.6% to 99.4%, with an overall transmission rate of 94.2% (Figure 2). These results strongly demonstrate that the *Medea* element described here can dominantly bias transmission in diverse *D. suzukii* populations.

**Figure 2.**
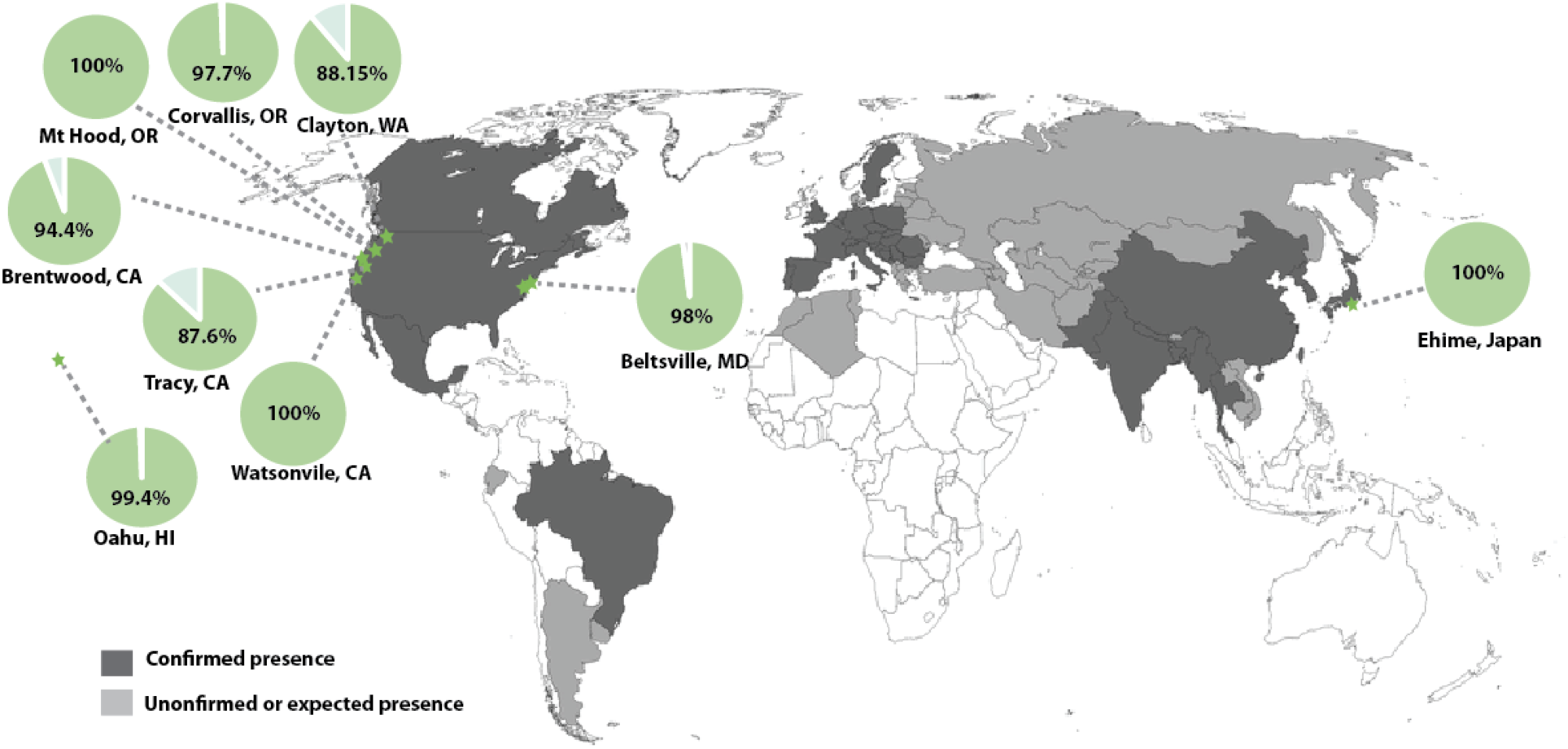
*Medea* functions in diverse populations of *D. suzukii*. Heterozygous *Medea/+* individuals were crossed with eight geographically distinct *D. suzukii* populations and *Medea* inheritance was measured. Overall, *Medea* biased inheritance with rates ranging from 87.6- 100%, suggesting that a *Medea* system generated in the laboratory could be utilized to manipulate some, but not all, diverse wild populations of *D. suzukii*. Green stars indicate the collection locations of the flies tested, green pie charts indicate the percentage *Medea* inheritance observed from heterozygous *Medea/+* females, and shaded areas on the map indicate locations where *D. suzukii* populations have been confirmed. The Corvallis, OR, strain was our reference *D. suzukii* strain used to engineer the *Medea*.

### Long Term Population Cage Experiments

The above observations suggested that *D. suzukii Medea* should be able to drive robust population replacement. To test this prediction, we mated *Medea*-bearing fathers to wildtype Corvallis, OR, strain mothers at three distinct introduction frequencies: low frequency (equal numbers of heterozygous *Medea*/+ and wildtype +/+ fathers mated to wildtype +/+ mothers); medium frequency (equal numbers of heterozygous *Medea*/+ fathers to wildtype +/+ mothers); and high frequency (equal numbers of homozygous *Medea*/*Medea* fathers to wildtype +/+ mothers). These experiments were conducted in separate bottles in biological triplicate for the low and medium threshold and quadruplicate for the high threshold drives, producing ten distinct populations with initial allele frequencies ranging from ∼12.5-50%. Altogether, these population cage experiments were followed for 19 generations, counting the number of *Medea-*bearing adults each generation, as described previously ^30,32^. Interestingly, the observed changes in *Medea* frequency over time indicated that, for release proportions of 50% or smaller, the *D. suzukii Medea* element was unable to drive into the wildtype population, likely because of selected drive resistance combined with high fitness costs outweighing the effect of drive.However, at higher introduction frequencies of >90%, the drive largely compensated for the fitness cost, allowing the construct to remain in the population at high frequencies for the duration of the experiment (19 generations; Figure 3).

**Figure 3.**
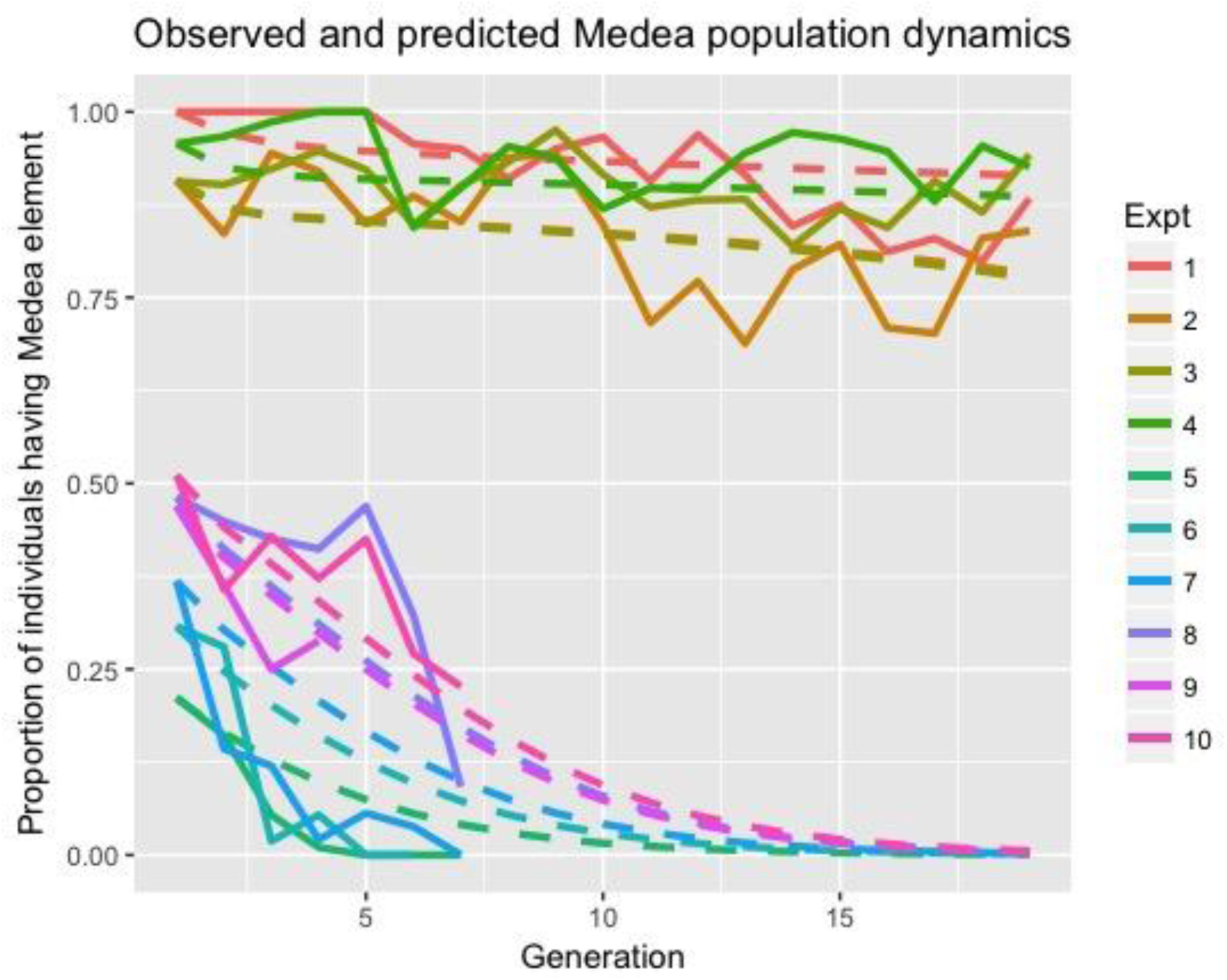
Observed and predicted dynamics of *D. suzukii Medea* element. Population cage experiments were set up by mating wild type (+/+) and homozygous *Medea* males (*Medea*/*Medea*) with wild type (+/+) females, producing a frequency of heterozygotes (*Medea*/+) in the first generation of 21-100%. Population counts were monitored over 19 generations. Results from these experiments are shown as solid lines, with fitted model predictions shown as dashed lines. Observed data are consistent with a toxin efficiency of 100% in *Medea*-susceptible mothers, 93% in *Medea*-resistant mothers (95% CrI: 90-95%), a heterozygote fitness cost of 28% (95% CrI: 27-30%), a homozygote fitness cost of 65% (95% CrI: 62-67%), and an initial resistant allele population frequency of 78% (95% CrI: 57-97%). For high initial heterozygote frequencies (90-100%), the element is capable of manipulating inheritance in its favor in order to maintain its presence at high population frequencies, despite a fitness cost. For lower initial heterozygote frequencies (∼50% or less), the element is eliminated from the population.

### Molecular Characterization of Resistance

To understand whether resistance of the target mRNA to the toxin played a role in observed *Medea* inheritance rates of <100%, we performed genomic DNA sequencing of the myd88 5’UTR miRNA target region from *Medea*/+ and +/+ progeny from the crosses described above. Genomic sequence analysis revealed that, out of 4 miRNA target sites, one to two sites were perfectly conserved in *Medea*/+ individuals (site #4 or sites #1 and #4, depending on the individual), while only one (site #4) was perfectly conserved in +/+ individuals (Supplementary Figure 1); additionally, for sites that were not perfectly conserved (i.e., had mutations; #1-3),some of the mutations were the same for *Medea*/+ and +/+ flies (for sites #2 and #3), while others were only found in one type of fly or the other (for site #1). To further this analysis, we also sequenced +/+ individuals from all of the geographically distinct populations tested above, and discovered a similar trend - i.e., that only one of the four miRNA target sites was perfectly conserved (#4), two others (#2 and #3) had the same mutations in all strains (including the *Medea*/+ and +/+ individuals above), and a third site (#1) had variable mutations that appeared to correlate with *Medea* efficiency. Together, these observations indicate that the nature of mutations differed between backgrounds correlating with observed *Medea* inheritance rates, suggesting that the efficiency of the miRNA “toxin” is likely influenced by resistance alleles, which influence *Medea* transmission.

### Mathematical Modeling

To characterize the population dynamics observed in the above cage experiments, we fitted a mathematical model to the observed data in which the *Medea* element had an associated fitness cost in heterozygotes and homozygotes and there was a *Medea*-resistant allele present in the population that reduced toxin efficiency. For the fitted model, the *Medea* element was estimated to have a toxin efficiency of 93% in individuals homozygous for the resistant allele (95% credible interval (CrI): 90-95%) and was assumed to have a toxin efficiency of 100% in individuals lacking the resistant allele. The *Medea* element was estimated to confer a large fitness cost on its host - 28% in heterozygotes (95% CrI: 27-30%) and 65% in homozygotes (95% CrI: 62-67%) - and the resistant allele was estimated to have an initial allele frequency of 78% in the population (95% CrI: 57-97%).

Predictive mathematical modeling based on these parameter estimates suggests that the *Medea* element would spread to fixation in the absence of toxin resistance if released above a threshold frequency of 79% (Figure 4A). Spread to fixation would also be expected if the fitness costs of the generated *Medea* element were halved (Figure 4C), even if all individuals in the population were homozygous for the *Medea*-resistant allele (Figure 4D), provided the element was released above a threshold frequency of ∼25-27%. Consistent with the experimental results (Figure 3), a *Medea* element with a large fitness cost in a *Medea*-resistant population is expected to be maintained at high frequencies through its drive; however, its eventual elimination is inevitable unless supplemental releases are carried out. However, for high release frequencies (90-95%), the element may be maintained at high frequencies (>75%) for ∼20 generations (Figure 4B), which likely exceeds the duration required for agricultural impact. Of note, the ability of the drive to counteract large fitness costs is significant, as demonstrated by comparison to non-driving alleles with analogous fitness costs that rapidly decline in frequency following a 95% release (black lines in Figures 4A and 4C).

**Figure 4.**
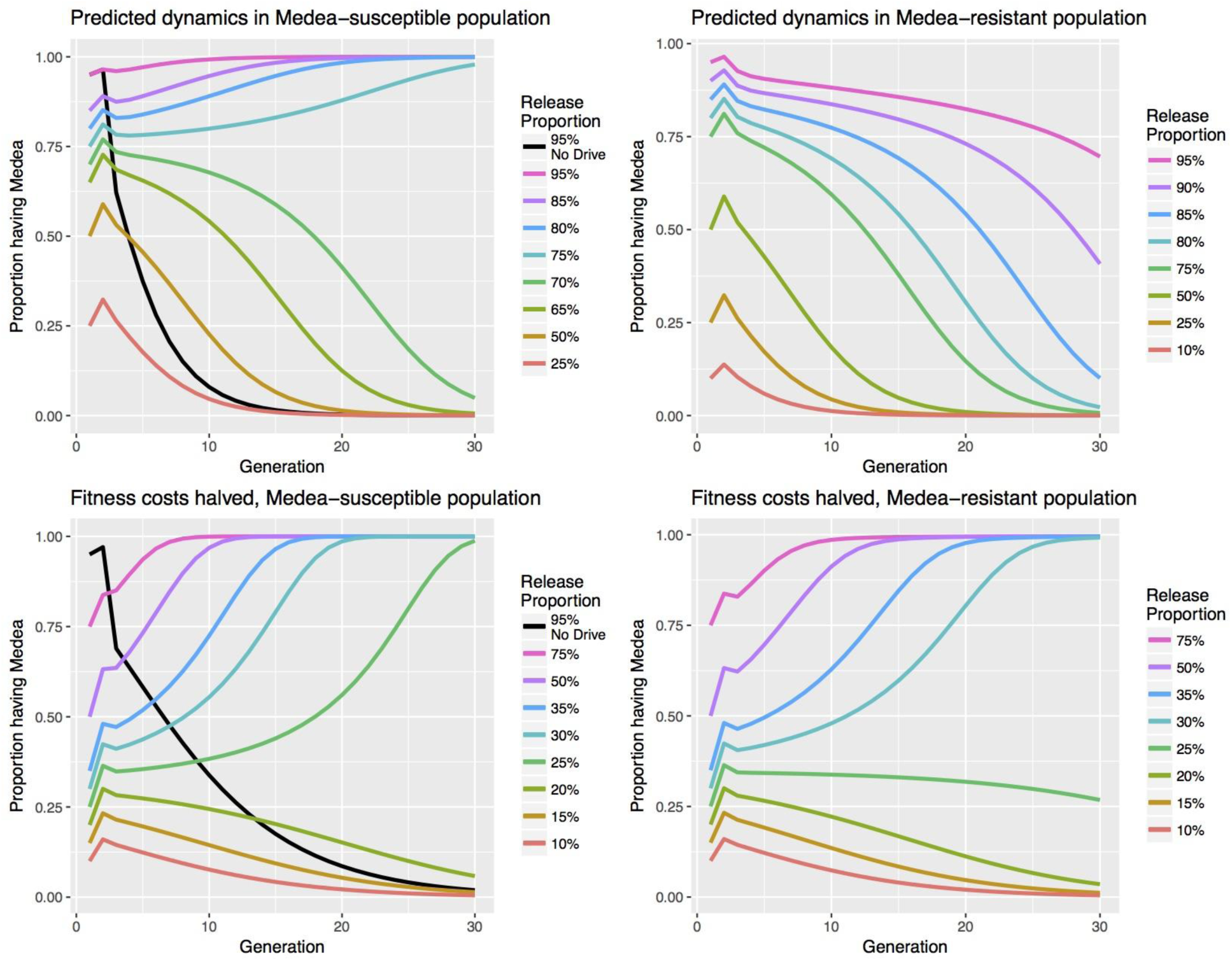
Predicted dynamics of *D. suzukii Medea* element. In all cases, releases of homozygous *Medea* males are assumed, except for black lines, which describe the dynamics of an equivalent release of non-*Medea* males. (A) In a *Medea*-susceptible population, the generated element (toxin efficiency = 100%, heterozygote fitness cost = 28%, homozygote fitness cost = 65%) displays threshold dynamics, spreading to fixation for release proportions of 73% or higher. (B) In a *Medea*-resistant population in which toxin efficiency is 93%, as inferred from the laboratory studies, the *Medea* element can be maintained at high frequencies following a high release proportion; however, its eventual elimination is inevitable unless supplemental releases are carried out. (C) In a *Medea*-susceptible population, if fitness costs are halved, the element displays threshold dynamics, spreading to fixation for release proportions of 23% or higher. (D) In a *Medea*-resistant population, if fitness costs are halved, the critical release threshold is raised slightly to 25%.

## Discussion

This study represents the first comprehensive characterization of a fully functional *Medea-*based gene drive element being challenged with pre-existing resistance in a long term multi-generational population cage experiment (19 generations). The synthetic *Medea* element described here showed maximal levels of dominance, up to 100% in some populations, but <100% inheritance bias in other populations. We hypothesized that this difference could be attributed to the presence of resistance arising from naturally occurring genetic variation that rendered certain embryos immune to the miRNA toxin. This hypothesis is supported by the sequencing data, as many of the sequenced miRNA target sites contained mutations that likely affected miRNA function and lowered toxin efficiency. Although we did not attempt to measure individual miRNA efficiency, it is possible that not all of the miRNAs are effective at target gene knockdown, and that particular target site mutations reduce toxin efficiency significantly enough to allow survival of some wildtype individuals. This is supported by sequencing data collected from the eight distinct geographic populations, which suggests that certain target site mutations are correlated with incomplete *Medea* dominance patterns.

The above observations highlight the importance of resistance as a possible impediment to the use of gene drives, including toxin-antidote drive systems, in the field ^18,26,28^. Multiple recent studies have highlighted resistance as a major obstacle to gene drive, mostly in the context of homing-based CRISPR/Cas9 drives ^39–43^. Although a *Medea* system may be less prone to resistance-associated spread impediment because, unlike homing-based drive, its mechanism of action is not likely to generate resistant alleles ^18,40^, it will face pre-existing resistant alleles given the natural genetic diversity found in wild populations. Furthermore, such mutations would be expected to face strong positive selection and increase in frequency over time, which would likely expand their effect. Therefore, any meaningful attempt at generating a *Medea-*based gene drive system capable of manipulating diverse wild populations must plan for, and mitigate the effects associated with, resistant alleles. This may be achieved in several ways. Firstly, sequencing-based characterization of naturally occurring genetic variation in geographically distinct target populations can help guide selection of target sites that are well conserved across all populations in which the drive is intended to function. Secondly, miRNA target site selection could be limited to the coding DNA sequence regions of a genome, which tend to be strongly conserved, as opposed to regions such as the 5’UTR, which canonically have higher tolerance for sequence variation. Thirdly, the choice of multiple target sites that have been validated to achieve knockdown and the creation of a polycistronic “toxin” can ensure that toxin efficiency is maximally high.

That said, modeling results suggest that a *Medea* element having a high fitness cost and high (though imperfect) toxin efficiency may be capable of maintaining itself in a population for a period of several years following a series of large-scale releases of homozygous males. Either decreasing the fitness cost of the element or minimizing resistance to the toxin are expected to enable the element to spread to fixation above a release threshold of ∼25-79% (the lower bound corresponds to halved fitness costs). While the stated release thresholds are high, they may be achievable given multiple successive releases (releases associated with the classical sterile insect technique for the Mediterranean fruit fly were of an even higher magnitude ^44^), and may be desirable for biosafety considerations related to novel genetic control strategies.

## Materials and Methods

### miRNA Design and Assembly

The *D. melanogaster* miRNA mir 6.1 stem-loop was modified to target four sites in the *Drosophila suzukii* myd88 5’UTR as previously described ^35,45^. The *Drosophila suzukii* myd88 gene ortholog was identified using the Augustus gene prediction tool ^46^, and sites were selected in the region spanning the ∼300 base pairs upstream of the start codon, which was presumed to be the 5’UTR. To generate mir6.1 stem-loop backbones that create mature miRNAs complementary to each of these target sites, pairs of primers were annealed and products were utilized for two subsequent rounds of PCR and cloned into the pFusA backbone (from the Golden Gate TALEN and TAL Effector Kit 2.0, Addgene cat#1000000024) using Golden Gate assembly ^47^ to generate plasmid OA-961C. Assembled miRNAs were then subcloned into final plasmid OA-961B using PacI and FseI. The *Drosophila suzukii* myd88 5’ UTR region with target sequences, and sequences of primers used in the miRNA cloning, are listed in Tables S1 and S2, respectively.

### Construct Assembly

To generate plasmid OA-961B, components were cloned into the *piggyBac* plasmid pBac[3xP3- EGFP afm] ^48^ using Gibson assembly/EA cloning ^49^. Specifically, the predicted *D. suzukii bottleneck* (bnk) promoter was amplified from *D. suzukii* genomic DNA using primers 961B.5 and 961B.6, the predicted *D. suzukii* myd88 coding region was amplified from *D. suzukii* genomic DNA using primers 961B.3 and 961B.4, and the SV40 3’UTR fragment was amplified from template pWalium20-10XUAS-3XFLAG-dCas9-VPR (addgene plasmid #78897) using primers 961B.1 and 961B.2. The pBac[3xP3-EGFP afm] plasmid was digested with AscI and FseI, and the above three fragments were cloned in via EA cloning. The resulting plasmid was then digested with PmeI, and the following fragments were cloned in via EA cloning: the predicted *D. suzukii* Bicaudal-C (BicC) promoter region amplified with primers 961B.7 and 961B.8 from *D. suzukii* genomic DNA, the SV40 3’UTR fragment amplified with primers 961B.9 and 961B.10 from template pWalium20-10XUAS-3XFLAG-dCas9-VPR (addgene plasmid #78897), the hr5-IE1 promoter region ^38^ amplified from vector pIEx-4 (Novagen plasmid #71235-3) using primers 961B.11 and 961B.12, and the dsRed-SV40 3’UTR fragment amplified from template pScarlessHD-DsRed (addgene plasmid #64703) with primers 961B.13 and 961B.14. Assembled miRNAs were then subcloned into final plasmid OA-961B using PacI and FseI. A list of primer sequences used in the above construct assembly can be found in Table S3; the *D. suzukii* myd88 coding region sequence, bnk promoter region sequence, and BicC promoter region sequence can be found in Table S1. The *Drosophila suzukii* myd88, BicC, and bnk gene orthologs were identified using the Augustus gene prediction tool ^46^

### Fly Culture and Strains

*Drosophila suzukii* wild type flies from Corvallis, Oregon, were a kind gift of P. Shearer, and were maintained under 12L:12D conditions at 20°C and 50% ambient humidity on a modified cornmeal medium, the recipe for which was obtained from the A. Ray Lab at UC Riverside. Rainbow Transgenics (Camarillo, CA) carried out the injections of construct OA-961B: the construct, along with a source of transposase, was injected into *D. suzukii* embryos using standard *D. melanogaster* injection procedures, and the surviving G_0_ adults were individually outcrossed to wildtype (WT) individuals. G_1_ progeny were screened for the presence of the *Medea* element (as evidenced by ubiquitous dsRed expression and eye-specific eGFP expression), and one G_1_ transformant male was recovered and outcrossed to wildtype *D. suzukii* females.

### *Medea* Genetic Behavior

To determine the genetic behavior of the *Medea* element in *D. suzukii* flies, male and female G_2_ progeny from the single obtained G_1_ transgenic male were individually outcrossed to wildtype (non-*Medea* bearing, +/+) *D. suzukii* flies, and the resulting progeny were scored for the presence of the *Medea* element; this process was repeated until the G6 generation, and the resulting data are presented in Table 1. To assess whether the *Medea* system would function well in geographically distinct populations of *D. suzukii*, heterozygous *Medea*/+ males were individually introgressed with virgin females from eight geographically distinct *D. suzukii* populations, and resulting heterozygous *Medea*/+ virgins were crossed back to males from the eight populations to determine whether the *Medea* element functioned as expected (Figure 2). A total of 1319 progeny were counted, and 96.4% (50% expected given traditional Mendelian inheritance) were *Medea*-bearing.

### Mathematical Modeling

Model fitting was carried out using Bayesian Markov Chain Monte Carlo methods in which parameters describing the population dynamics of the *Medea* element were estimated, including 95% CrIs. Estimated parameters include fitness costs associated with being heterozygous, *s*_*Het*_, or homozygous, *s*_*Hom*_, for the *Medea* element, and the reduced maternal toxin efficiency associated with the *Medea*-resistant allele, *e*_*R*_, present in the population at a given initial frequency, *p*_*R*_. Prior information on the parameter *e*_*R*_ was inferred from G_5_ and G_6_ outcrosses in which heterozygous *Medea* females were mated with wildtype males and the proportion of wildtype offspring was nonzero. A simplified version of the fitted model was used to infer the expected dynamics of the generated *Medea* element and one with its fitness costs halved in both a fully *Medea*-susceptible population and a fully *Medea*-resistant population. The modeling framework is described in the Supplementary Material.

### Embryo Viability Determination

For embryo viability counts (Table 2), adult virgin females were mated with males of the relevant genotypes for 2-3 days in egg collection chambers with plates containing modified cornmeal medium, supplemented with dry yeast. Then, an overnight egg collection was carried out, after first having cleared old eggs from the females through a pre-collection period on a separate plate for several hours. All embryos (between 165-301) were counted, and kept on an agar surface at 20°C for 48 hours. The % survival was then determined by counting the number of unhatched embryos. Each experiment was carried out in duplicate, and the results presented are averages from these two experiments. Embryo survival was normalized with respect to the % survival observed in parallel experiments carried out with wild type flies, which was 91.63±0.26%. For adult fly counts (Table 2), the adult flies used for each embryo count assay replicate were transferred from egg collection plates to 250 ml bottles containing modified cornmeal medium, allowed to lay eggs for 24 hours, and then removed. 100% of the resulting progeny (between 116-206 progeny) from these bottles were counted, and the results of the two replicates for each experiment were averaged together.

### Population Cage Experiments

To determine whether the generated *D. suzukii Medea* is capable of spreading through populations, population cage experiments were set up as follows. Heterozygous *Medea*/+ males and virgin females were crossed to each other for multiple generations to generate homozygous stocks; homozygosity was confirmed by outcrossing. Then three types (low, medium, and high threshold, the first two in triplicate and the last in quadriplicate) of drive experiments were set up by crossing *Medea*-bearing males with wildtype flies of the strain from Corvallis, OR, of a similar age in 250ml bottles containing modified cornmeal medium. Low frequency drives had equal numbers of heterozygous *Medea*/+ and wildtype +/+ males mated to wildtype +/+ virgins, *Medea* allele frequency of ∼12.5%; medium frequency drives had equal numbers of heterozygous *Medea*/+ males mated to wildtype +/+ virgins, *Medea* allele frequency of ∼25%; and high frequency drives had equal numbers of homozygous *Medea*/*Medea* males mated to wildtype +/+ virgins, *Medea* allele frequency of ∼50%. The total number of flies for each starting population was 100. After being placed together, adult flies were removed after nine days. After another seven days, half of the progeny (randomly selected) were counted, and the other half were placed in a new bottle to continue the simulation, and this process continued throughout the duration of the experiment. All fly experiments were carried out in the conditions described above.

### Target Site Genotype Screening

To identify the genotypes of the four miRNA target sites of various flies from the *Medea* outcrosses and drive experiment and of different *D. suzukii* strains, genomic DNA was extracted from individual flies with the DNeasy Blood & Tissue kit (QIAGEN) following the manufacturer’s protocol. PCR was conducted using standard procedures to amplify the target loci using primers 807G and 807H (Supplementary Table 2), which amplified a region of ∼550 base pairs in the myd88 5’UTR region. The PCR program utilized was as follows: 98°C for 30 seconds; 35 cycles of 98°C for 10 seconds, 57°C for 20 seconds, and 72°C for 30 seconds; then 72°C for 10 minutes. The PCR products were purified with the MinElute^?^ PCR Purification Kit (QIAGEN) according to the manufacturer’s protocol and sent for Sanger sequencing (SourceBioScience) using the same primers as utilized for the PCR. Target site sequences were analyzed and aligned with DNAStar software.

## Author Contribution(s)

A.B. and O.S.A conceived and designed experiments. A.B., D.O, T.Y. performed all molecular and genetic experiments. J.M.M performed mathematical modelling. A.B., J.M.M., and O.S.A. contributed to the writing of the manuscript. All authors analyzed the data and approved final manuscript.

## Acknowledgments

This work was supported by a grant from the California Cherry board awarded to O.S.A. We thank Joanna Chiu (UCD), Anandasankar Ray (UCR) and Peter Shearer (Oregon State) for providing the various *D. suzukii* strains used in this study.

---

**Supplementary Figure 1.**
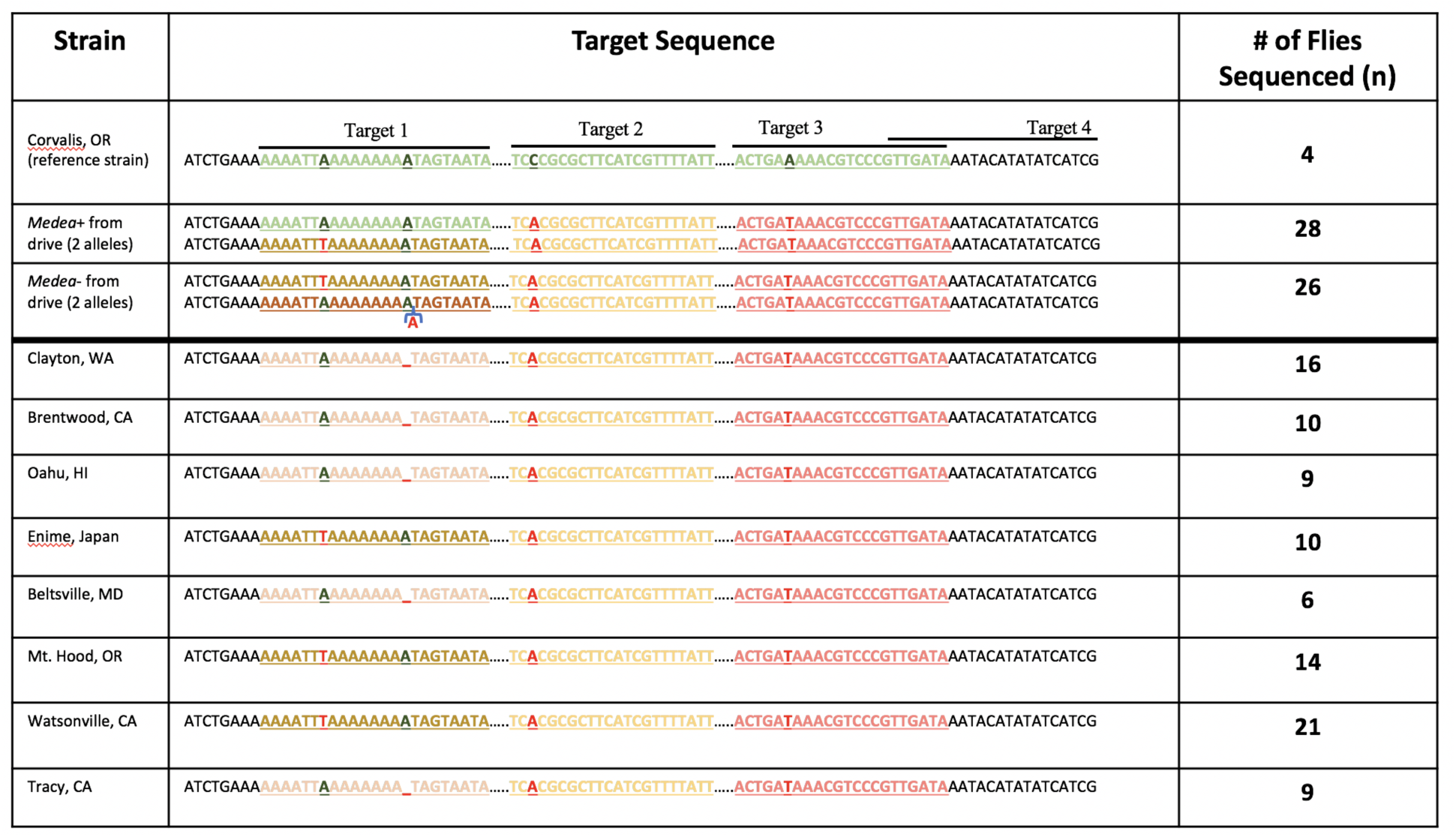
Genomic DNA sequences of the myd88 5’UTR region of various strains/fly types targeted by the *Medea* toxin miRNAs. Green and black nucleotides represent sequence perfectly complementary to the miRNAs; red and other color nucleotides represent specific mutations and target sites that are not perfectly complementary to the miRNAs, respectively. Target site four is not highlighted/underlined as it is perfectly conserved among all sequenced flies.

**Supplementary Table 1.**
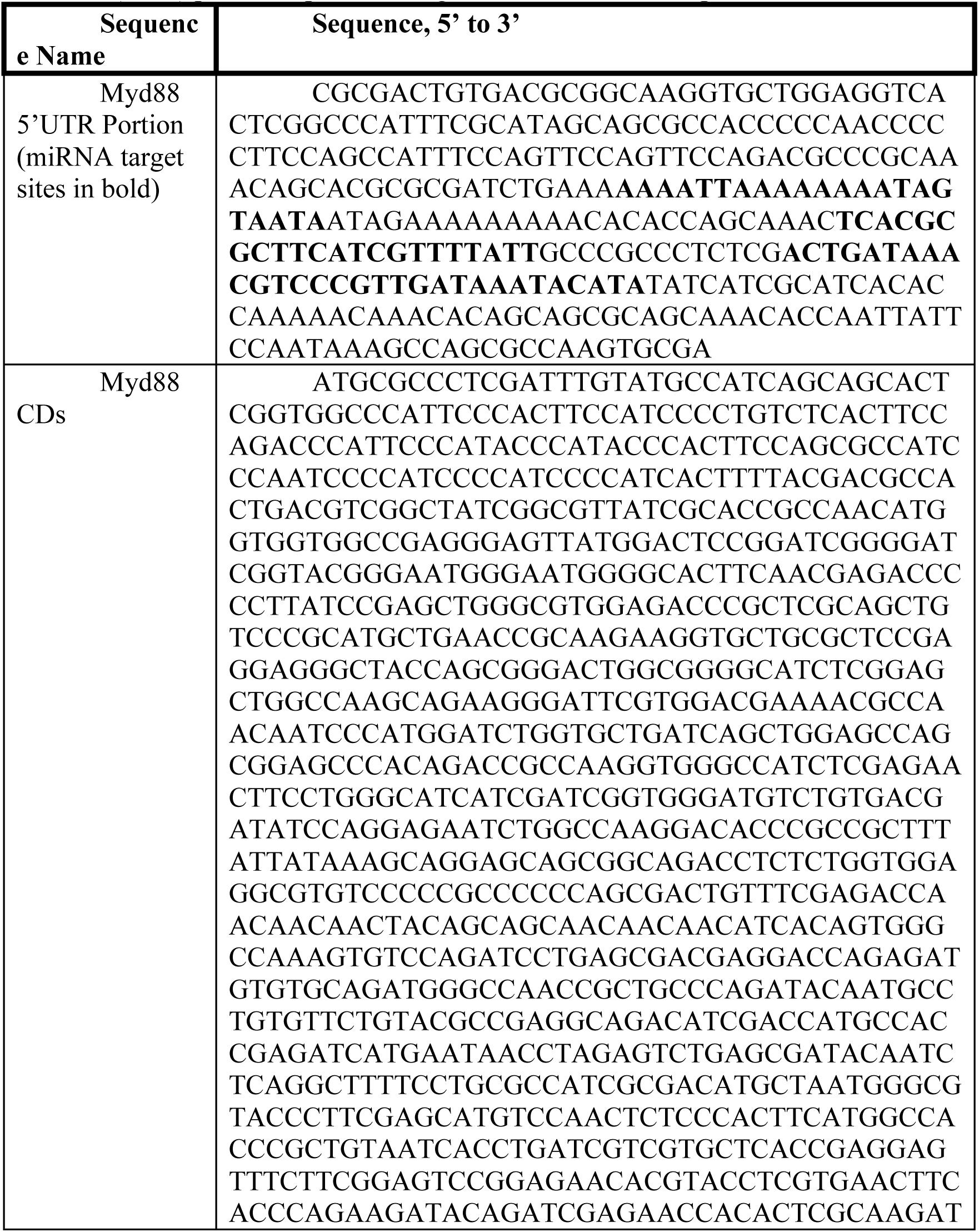
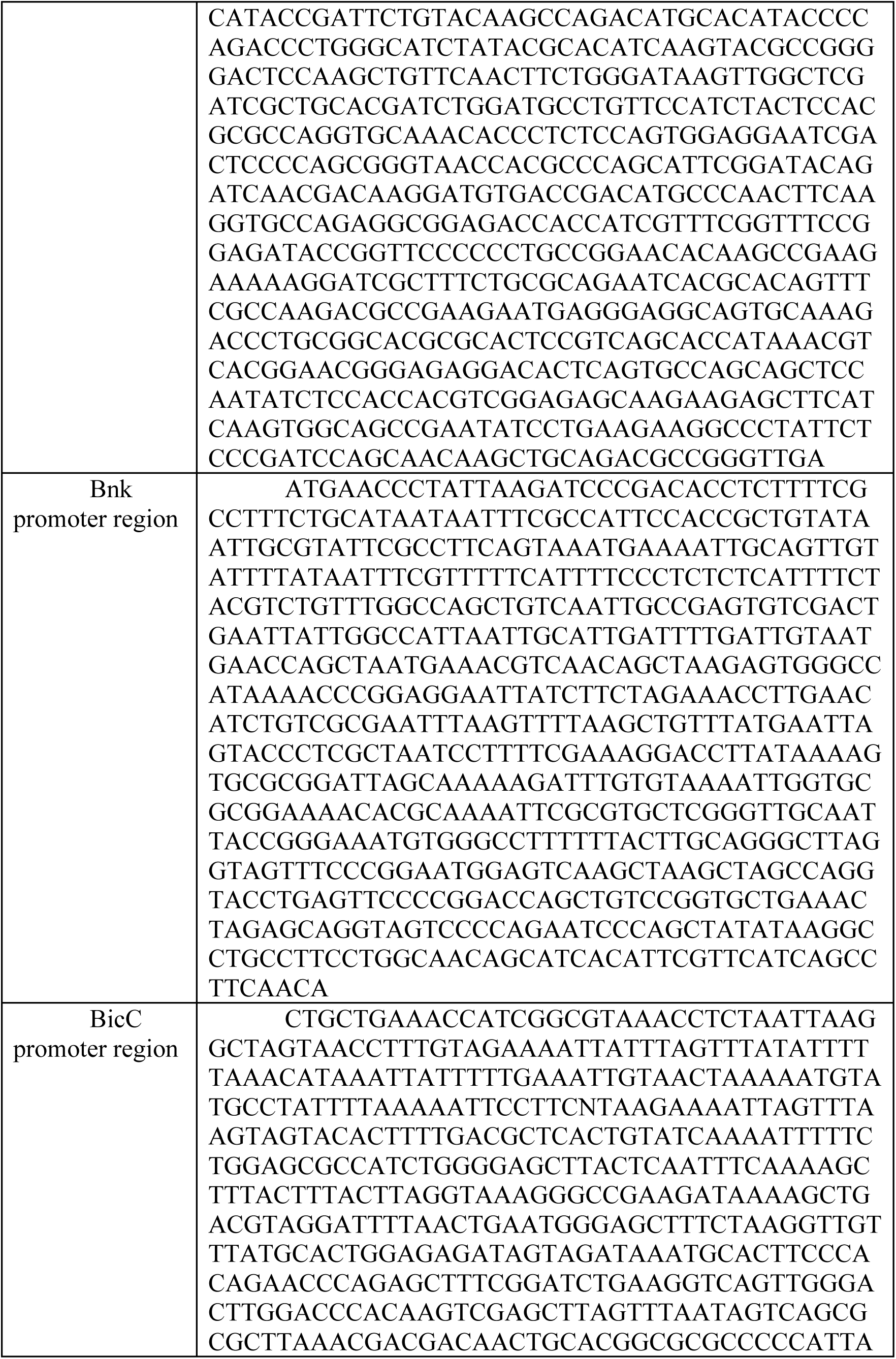
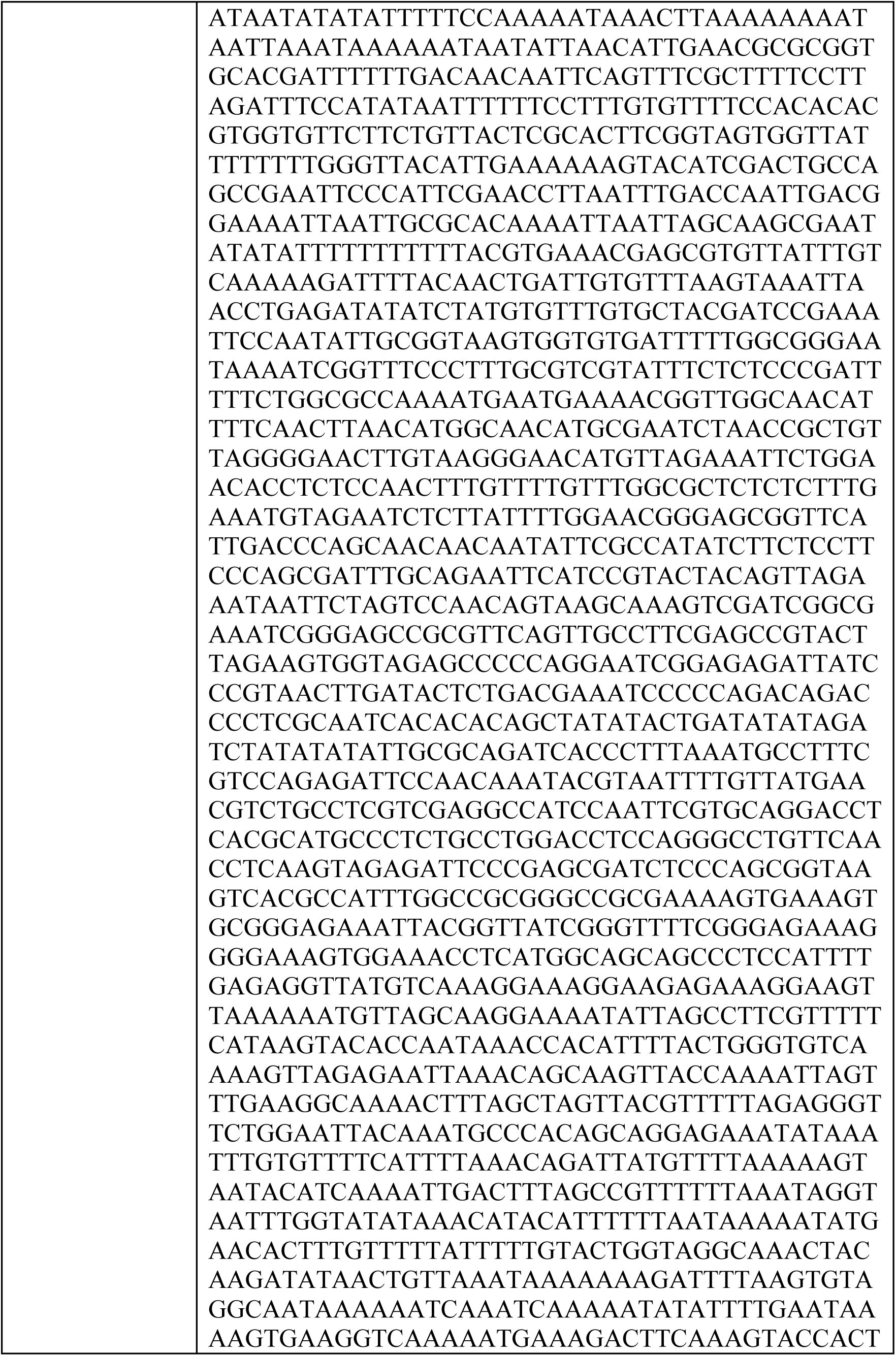
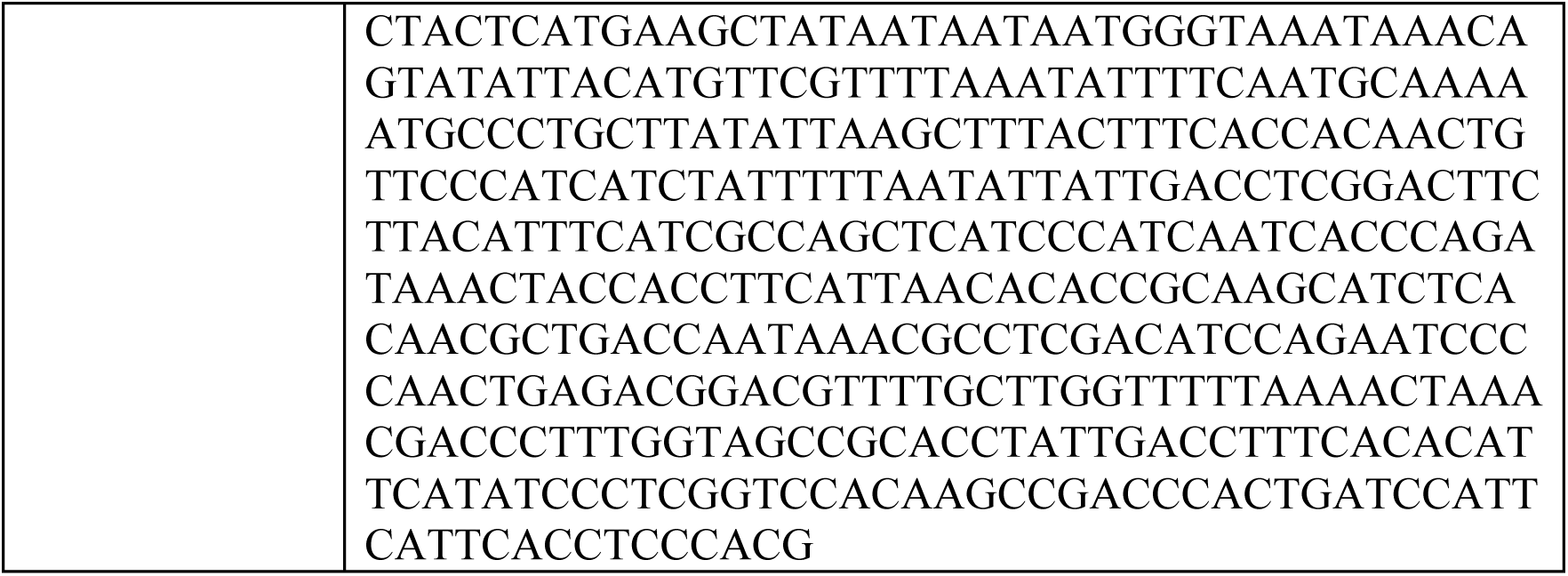
*D. suzukii* genomic DNA sequences used in constructing synthetic *Medea*. Genomic DNA sequences of: the targeted portion of the myd88 5’UTR, with miRNA target sites indicated in bold (site three and four overlap); the myd88 CDs used as the rescue gene; the bottleneck (bnk) predicted promoter region utilized to drive expression of the rescue; and the bicoid (BicC) predicted promoter region utilized to drive expression of the rescue.

**Supplementary Table 2.**
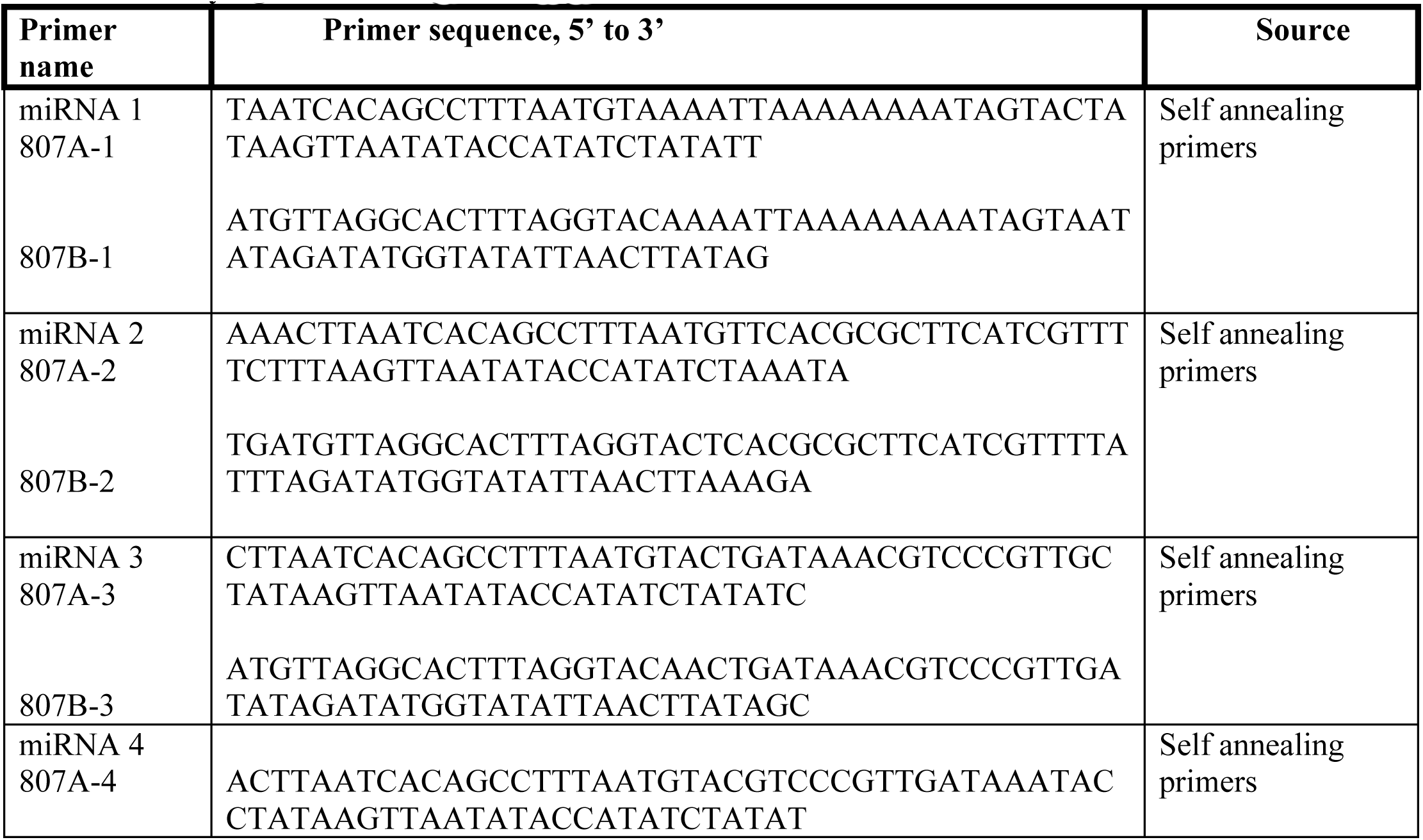
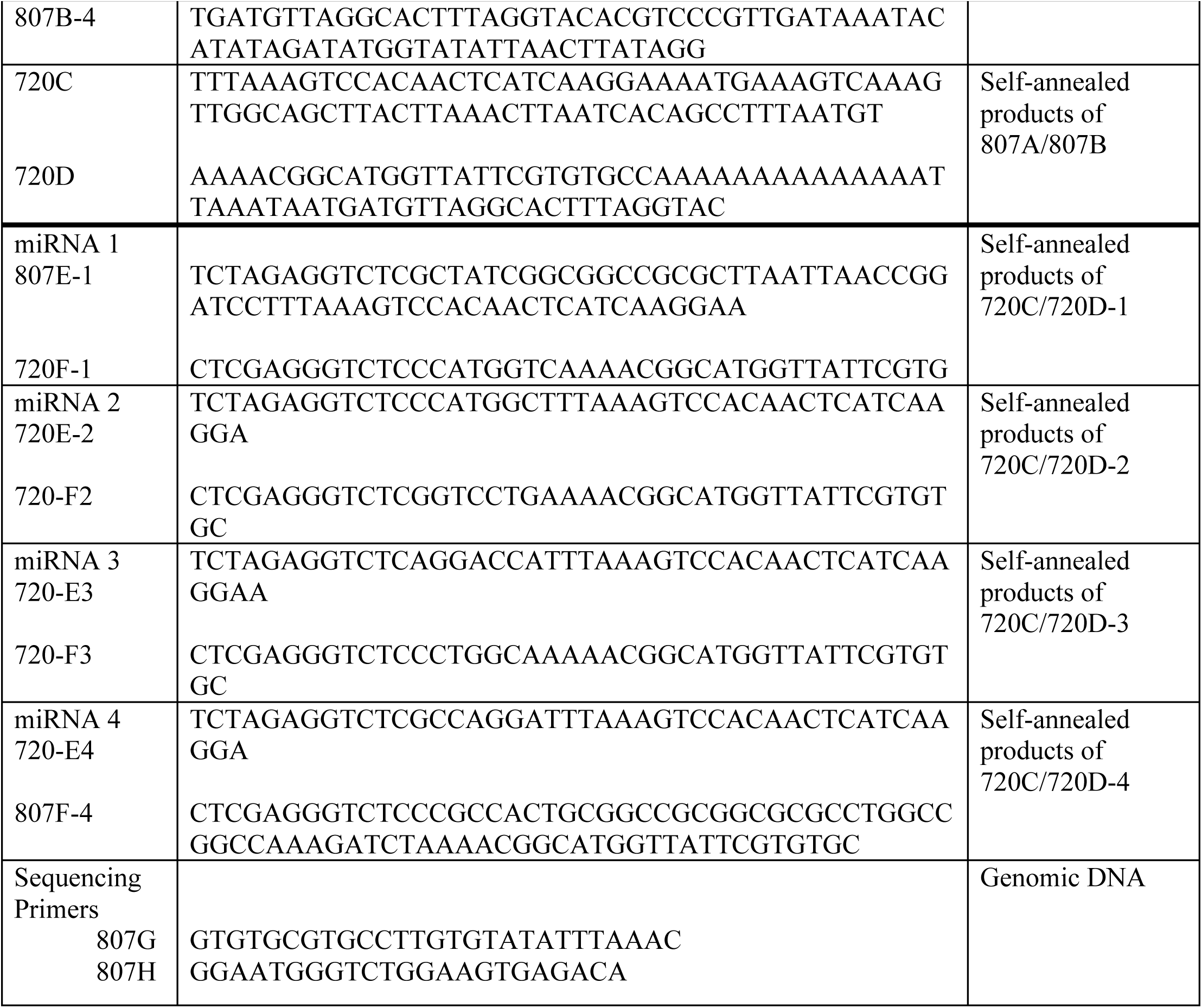
Sequences of primers utilized in cloning myd88-targeting miRNA and verifying miRNA target site regions.

**Supplementary Table 3.**
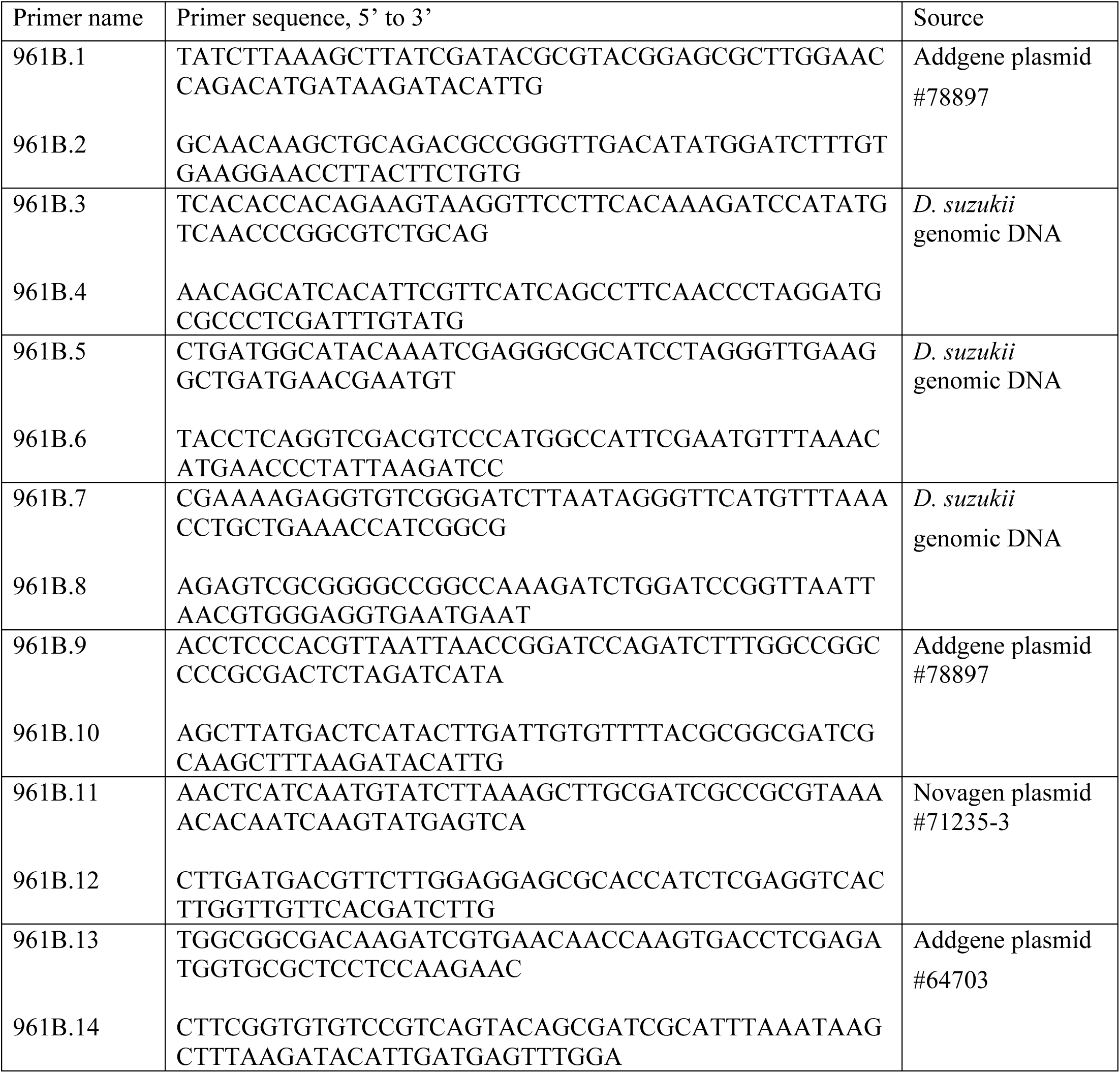
Sequences of primers used to assemble plasmid OA-961B.

## Supplemental Methods

### Model Fitting

We modeled *Medea* dynamics under laboratory cage conditions assuming random mating and discrete generations. To model resistance to the maternal toxin, we considered a *Medea* allele, “M”, and an unlinked toxin-resistant allele, “R”, that diminishes toxin efficacy in mothers having at least one copy of the *Medea* allele. We denote the absence of the *Medea* allele by “m” and the absence of the toxin-resistant allele by “r”. Consequently, there are nine possible genotypes – MMRR, MMRr, MMrr, MmRR, MmRr, Mmrr, mmRR, mmRr and mmrr. We denote the proportion of organisms having each genotype at the *k*th generation by 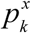, where *x* denotes one of the nine genotypes.

Given the large number of possible mating pairs (81), it is not feasible to show the complete equations for the next generation genotype frequencies here, so we instead describe them in brief. We considered the M and R loci as being independently inherited and following Mendelian inheritance rules, with the exception that mm offspring of mothers heterozygous for the *Medea* allele have reduced viability. If a *Medea*-heterozygous mother does not have any copies of the resistant allele (i.e. has the genotype Mmrr), the viability of their mm offspring is reduced by 100%; however if the mother has two copies of the resistant allele (i.e. has the genotype MmRR), then the viability of their mm offspring is reduced by a fraction, *e*_*R*_, denoting the maternal toxin efficiency in the presence of the resistant allele. We considered two models for maternal toxin efficiency in MmRr mothers – in model A, the toxin efficiency is *e*_*R*_, and in model B, the toxin efficiency is (1+*e*_*R*_)/2, i.e. midway between that of MmRR and Mmrr mothers.

These considerations allow us to calculate the expected genotype frequencies in the next generation before accounting for fitness costs. Let us denote these frequencies by **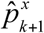**, where *x* denotes the genotype and *k*+1 denotes the next generation. Normalizing these ratios to account for a fitness cost, *s*_*Het*_, associated with being heterozygous for the *Medea* allele, and *s*_*Hom*_, associated with being homozygous for the *Medea* allele, the genotype frequencies in the next generation are given by:

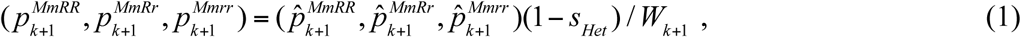

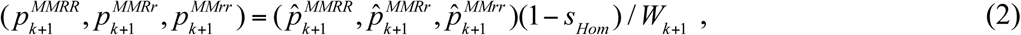

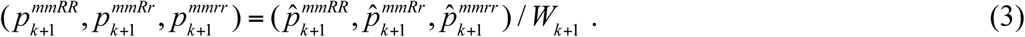

Here, *W*_*k*+1_ is a normalizing term given by,

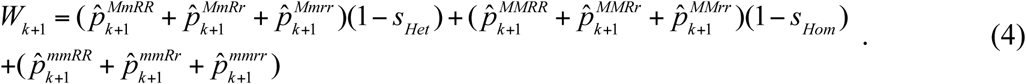

The likelihood of the population cage data was calculated by assuming a binomial distribution of wildtype and *Medea*-bearing individuals, and by using the model predictions to generate expected proportions for each set of parameter values, i.e. by calculating the log-likelihood,

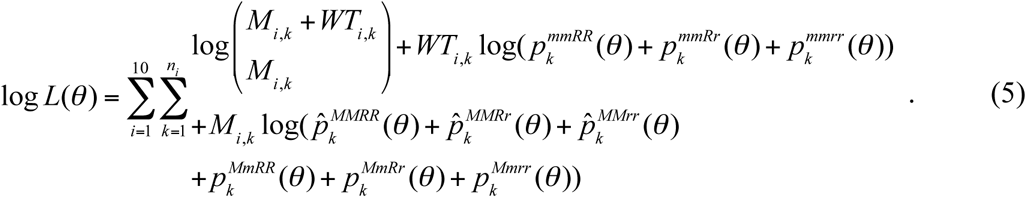

Here, *M*_*i, k*_ and *WT*_*i, k*_ are the number of *Medea*-bearing and wildtype individuals at generation *k* inexperiment *i*, the *i*th experiment is run for *n*_*i*_ generations, and the expected genotype frequencies are dependent on the model parameters *θ* ={*e*_*R*_, *p*_*R*_, *s*_*Het*_, *s*_*Hom*_}, where *e*_*R*_, *s*_*Het*_ and *s*_*Hom*_ are as defined earlier and *p*_*R*_ is the resistant allele frequency at the beginning of the experiment. The initial condition for each experiment was such that, for the *Medea* allele, heterozygote frequency was determined according to the first generation data with the remaining individuals being wildtype, and for the resistant allele, Hardy-Weinberg equilibrium was assumed with a resistant allele frequency of *p*_*R*_ and the resistant allele being independently distributed from the *Medea* allele.

Prior information on *Medea* toxin efficiency in *Medea*-resistant mothers was inferred from G_5_ and G_6_ outcrosses in which heterozygous *Medea* females were mated with wildtype males and the proportion of wildtype offspring, *p*(WT), was nonzero. In *Medea*-resistant mothers, for a toxin efficiency of *e*_*R*_, the ratio of wildtype to *Medea*-bearing offspring should be (1 − *e*_*R*_): 1, and hence the proportion of wildtype offspring should be (1– *e*_*R*_) / (1 + 1– *e*_*R*_). Rearranging this equation, then toxin efficiency in terms of _*p*_(WT) is given by *eR* = (1– 2 _*p*_(*WT*)) / (1– _*p*_(*WT*)). Using this equation, we calculated the mean and variance of toxin efficiency in *Medea*-resistant mothers from the results of 17 G_5_ and G_6_ outcrosses displaying resistance, from which we parameterized a normally distributed prior on *e*_*R*_.

We used a Bayesian Markov Chain Monte Carlo (MCMC) sampling procedure to estimate our model parameters, including 95% credible intervals, and used the Deviance Information Criterion (DIC) as a criterion to select between the two models for toxin efficiency in MmRr mothers. Following Gelman *et al.* (2004), we calculated the DIC as,

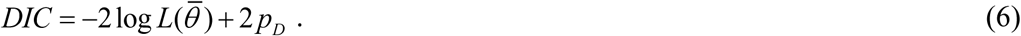

Here, *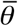* is the posterior mean of the model parameters, and *p*_*D*_ is the effective number of parameters as inferred from the MCMC chain, which may be calculated as,

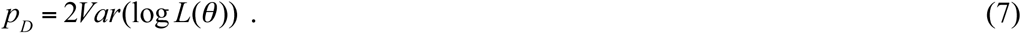

This favored model B, in which the toxin efficiency in MmRr mothers is midway between that of MmRR and Mmrr mothers (the model with smallest DIC value is favored and model A had a DIC value of 1146.5 while model B had a DIC value of 1145.3).

### Model Predictions

We modeled the expected dynamics of the generated *Medea* element and one with its fitness costs halved in both a fully *Medea*-susceptible population and a fully *Medea*-resistant population. Under these scenarios, a simpler model could be used since the resistant allele frequency is not expected to change. Assuming random mating and discrete generations, the proportions of the *k*th generation that are wildtype, heterozygous and homozygous for *Medea* are denoted by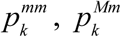 and 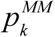, respectively.Considering all possible mating pairs, and taking into account that most wildtype offspring of heterozygous mothers are unviable – the viable fraction is denoted by (1-*e*), where *e* represents the maternal toxin efficiency – the genotypes of embryos in the next generation are described by the ratio 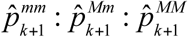, where,

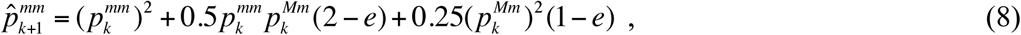

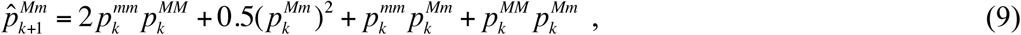

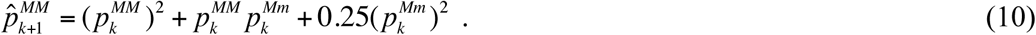

Toxin efficiency, *e*, equals 1 in a fully *Medea*-susceptible population, *e*_*R*_ in a fully *Medea*- resistant population, and 0 for a non-*Medea* allele. Normalizing these ratios and taking into account fitness costs, the genotype frequencies in the next generation are given by,

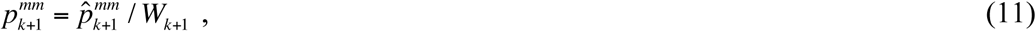

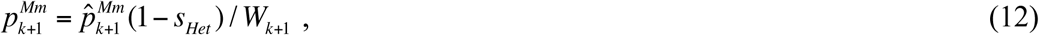

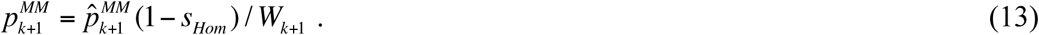

Here, *s*_*Het*_ and *s*_*Hom*_ represent the fitness costs associated with being heterozygous or homozygous for the *Medea* element, and *W*_*k*+1_ is a normalizing term given by,

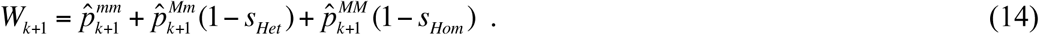

The only further change to the model used for projections is that release proportion refers to a release of homozygous *Medea* males, with the remainder of the first generation being half wildtype males and half wildtype females. Denoting the release proportion by *r*, then the *Medea* genotype frequencies in the second generation are given by:

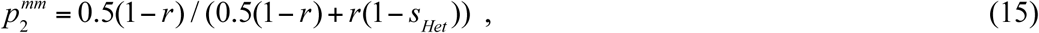

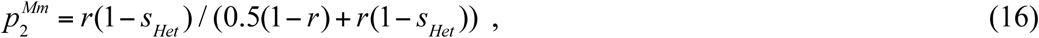

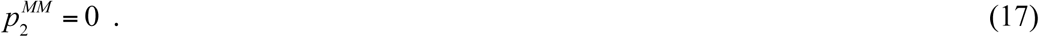

The model described in Equations 8-14 applies from generations 3 onwards.

